# Pseudopaline-mediated zinc uptake by *Pseudomonas aeruginosa* determines specific clinically relevant phenotypes and infection outcome

**DOI:** 10.1101/2025.01.08.631948

**Authors:** L Bosc, T Sécher, G Ball, D Le Pennec, M Tribout, M Ba, Y Bai, L Ouerdane, P Arnoux, Y Denis, X Lei, C Bordi, N Heuzé-Vourc’h, S Häussler, NO Gomez, R Voulhoux

**Author notes:** These authors contribute equally to the work (L Bosc, T Sécher).

## Abstract

The host-pathogen interface is a biological niche in which two entities competes for essential resources. The host’s nutritional immunity restrict access to metals, while a successful pathogen overcomes these restrictions using dedicated uptake pathways. Pseudopaline is a high-affinity metallophore allowing *Pseudomonas aeruginosa* to acquire zinc in chelated environments. We demonstrate that this pathway is the last-resort solution to acquire zinc for this dreadful pathogen. The capacity to provide this metal to zinc-metalloproteins drives clinically relevant phenotypes, such as the capacity to form a mature and antibiotic-tolerant biofilm, or to affect the outcome of an infection. These results place pseudopaline as a potential drug target for blocking *P. aeruginosa* pathogenic capacity and resensitizing established biofilm to classic antibiotic treatment.

**ABSTRACT:** Biological metals are essential trace elements which are required by metalloproteins, involved in virtually every cellular, structural and catalytic function of the bacterial cell. Bacterial pathogenesis involves a tug-of-war between the host nutritional immunity, sequestering essential metals and the invading pathogens that deploy high-metal affinity uptake strategies in order to overcome these defence mechanisms. Metallophores are high-affinity, low-molecular mass metal chelators produced and secreted by bacteria to access chelated metals from the environment. Pseudopaline is a metallophore produced and secreted by *Pseudomonas aeruginosa* to acquire zinc when the bioavailability of this metal is severely restricted, as in the presence of a strong metal chelator such as EDTA, or during infections when the nutritional immunity of the host is active, in mammals through the production of the zing binding protein calprotectin. We show that under the conditions of metal deprivation, a pseudopaline-deficient *P. aeruginosa* strain exhibit a severe intracellular zinc deficiency, establishing that the pseudopaline pathway is the last-resort and unique pathway for the bacteria to acquire zinc under these restricted growth conditions. The present study explores the pleiotropic role of pseudopaline-mediated zinc acquisition on several clinically relevant phenotypes and its capacity to drive infection outcomes, placing this machinery as a promising therapeutic target for *P. aeruginosa*’s infection, acting synergistically as a pathogenicity determinant as well as an adaptative trait allowing the establishment of a mature and antibiotic resistance biofilm necessary for recalcitrant chronic infections.

## INTRODUCTION

Biological metal ions are essential across all kingdoms of life (Nielsen, 2000). Their unique chemical features provide structural and catalytic properties to a wide-range of metalloproteins, broadly distributed in living organisms (Bertini et al., 2001). Zinc is, after iron, the second most abundant metal in living cells, involved in 6 to 9% of the prokaryotic and eukaryotic proteomes, respectively (Andreini et al., 2006; Andreini et al., 2009). In bacteria, zinc metalloproteins are involved in several functions essential to growth, such as central metabolism or DNA replication and repair, but also to virulence, antibiotic resistance, motility, extracellular protease activities, and modulation of the immune response (Mikhaylina et al., 2018; Palmer and Skaar, 2016) Biological metals are inorganic resources that need to be acquired by bacteria from the environment where they are often rare, either because of low concentration, restricted bioavailability, or competition between the inhabitants of the same ecological niche. In such competitions between pathogenic bacteria and host during bacterial infection, both protagonists have developed competing strategies for either sequestering or acquiring the much-coveted metal. Hence, the host synthetizes high-affinity chelators which withhold essential metals through the process referred to as “nutritional immunity” (Hood and Skaar, 2012). Conversely pathogens have evolved powerful metal uptake systems involving high affinity secreted molecules called metallophores (Cunrath et al., 2016) During bacterial infections, availability of zinc to pathogens is drastically reduced due to its sequestration by the host zinc-binding protein calprotectin (Hood and Skaar, 2012; Kehl-Fie and Skaar, 2010; Nelson et al., 2021; Vermilyea et al., 2021; Wakeman et al., 2016). Particularly, in cystic fibrosis (CF) patients the levels of calprotectin in bronchoalveolar lavage fluid (BAL) and serum are elevated and used as a biomarker of CF lung disease correlating with exacerbation (Gray et al., 2010; MacGregor et al., 2008; Wilson et al., 1975). To responsively overcome the growth-limiting zinc deficiency in tissues, imposed by the nutritional immunity, the human pathogen, *P. aeruginosa* has developed effective zinc import mechanisms judiciously de-repressed under intracellular zinc deficiency by the Zur regulator (formerly known as np20). Zur is a cytoplasmic protein, that prevents gene transcription by binding to the *zur* box of its promoter region and magnitude of repression is proportional to the intracellular zinc concentration (Patzer and Hantke, 1998). Thus, lower the zinc intracellular zinc concentration, the more active Zur is, and the greater the transcription of the initially repressed gene (Ellison et al., 2013). *P. aeruginosa* is therefore able to sense zinc scarcity in its environment in order to react by de-repressing the Zur regulon, comprising, among others, the Znu and Cnt (also called Zrm) zinc import mechanisms (Pederick et al., 2015; Vermilyea et al., 2021). The Znu pathway, widely distributed in bacteria, recognizes and imports free zinc in the cytoplasm via the periplasmic substrate-binding protein ZnuA and the inner membrane importer ZnuBC (D’Orazio et al., 2015; Pederick et al., 2015). In order to acquire zinc when it is not free but sequestered by a powerful chelator such as Ethylenediaminetetraacetic acid (EDTA) or calprotectin, *P. aeruginosa* relies on the dedicated Cnt pathway involving the high zinc affinity zincophore pseudopaline (Gomez et al., 2020; Lhospice et al., 2017; Mastropasqua et al., 2017; Zhang et al., 2019). Utilizing this mechanism, pseudopaline is first synthetized in the cytoplasm by CntL and CntM, then secreted in the milieu through the bacterial envelope by the CntI inner membrane exporter coupled to the outer membrane MexAB-OprM efflux pump. The extracellular zinc-bound pseudopaline is finally recovered by the bacteria through its specific outer membrane receptor, CntO. Under zinc sequestering conditions, the Cnt pathway appears to be the exclusive way for efficient zinc import from the medium since a *P. aeruginosa cntL* mutant - with a functional Znu pathway - is unable to maintain a normal intracellular zinc homeostasis when it is grown in zinc-deprived minimal chelating medium (MCM) consisting of minimal succinate (MS) medium supplemented with 100µM of the divalent metal chelator EDTA (Lhospice et al., 2017).

Numerous transcriptomic studies have reported a systematic up-regulation of the *cnt* operon under the infection conditions of CF patients or in medium reproducing these conditions (Bielecki et al., 2011; Cornforth et al., 2018; Kordes et al., 2019; Mastropasqua et al., 2018; Rossi et al., 2018; Son et al., 2007; Turner et al., 2015; Vermilyea et al., 2021). Together with the requirement of the Cnt pathway for *P. aeruginosa* growth in airway mucus secretion (Gi et al., 2015), these data demonstrate that the Cnt pathway is not only induced but also critical for the pathogen during infections.

In this study, we demonstrate the pleiotropic role of pseudopaline in *P. aeruginosa* infections. We first establish its essential role to secure the intracellular zinc concentration required for optimal planktonic growth of the pathogen in MCM medium. Moreover, we show that in such zinc limiting conditions, the Cnt pathway is necessary to establish a mature biofilm which provides protection from antibiotic exposure. We further explored the role of pseudopaline in environments encountered during *P*.

*aeruginosa* infection of the host. Specifically, we demonstrate that it plays a role in various steps in interactions with macrophages ranging from adherence and phagocytosis to intracellular survival and induction of immune responses. Additional *in vivo* experiments in a mouse model of *P. aeruginosa* lung infection confirmed the requirement for this zincophore to colonize the lung and develop a full virulent phenotype.

## RESULTS and DISCUSSION

### In the metal-sequestered growth condition, pseudopaline is responsible for maintaining intracellular zinc concentrations

Metals’ homeostasis in bacterial cells is at the centre of a tight regulation network, controlling both uptake pathways when facing scarcity or detoxification in case of metal excess. The growth in metal-restricted environments leads to the expression of specific and highly efficient metal import mechanisms to maintain intracellular metal homeostasis and ensure that metalloproteins maintain functional and operational requirements. This is the case for the metallophore pseudopaline whose production, secretion and recovery machineries encoded by the *cnt* operon are under the control of the Zur repressor and allows zinc acquisition even when the latter is chelated in the external environment. Zur repression is directly corelated to the intracellular zinc concentration since less Zur is loaded with zinc, less active it becomes in its ability to bind and repress promotors of *zur* regulon leading to higher the production of pseudopaline (Ellison et al., 2013). Under conditions of low zinc bioavailability due to extracellular sequestration by EDTA, as the one created in MCM (Lhospice et al., 2017), the normal intracellular zinc level and optimal planktonic bacterial growth are ensured by pseudopaline since in its absence (i.e. in the PA14Δ*cntL* mutant strain), the intracellular zinc content decreases and bacterial growth is impaired compared to the parental PA14 wild type (WT) strain (Fig. 1A and B and as previously shown (Gomez et al., 2020)). Further transcriptional experiments showing a similar *cntM* expression in MCM for both WT and Δ*cntL* PA14 strains are excluding any polar effect of *cntL* in frame deletion on the downstream *cntM* gene expression (Fig. S1). The growth defect of the *cntL* mutant depends exclusively on the presence of pseudopaline as exogeneous addition of 1μM of synthetic pseudopaline restores the growth defect of the PA14Δ*cntL* mutant (Fig. 1B, grey curve). This form of complementation of pseudopaline synthesis defect in the *cntL* mutant with externally-acquired pseudopaline not only exclude any polar effect of the deletion but also confirms that the mode of action of pseudopaline involves an extracellular acquisition stage, in agreement with the three proposed steps: synthesis, secretion and recovery of the pseudopaline-dependant zinc uptake pathway (Gomez et al., 2020).

**FIGURE 1.**
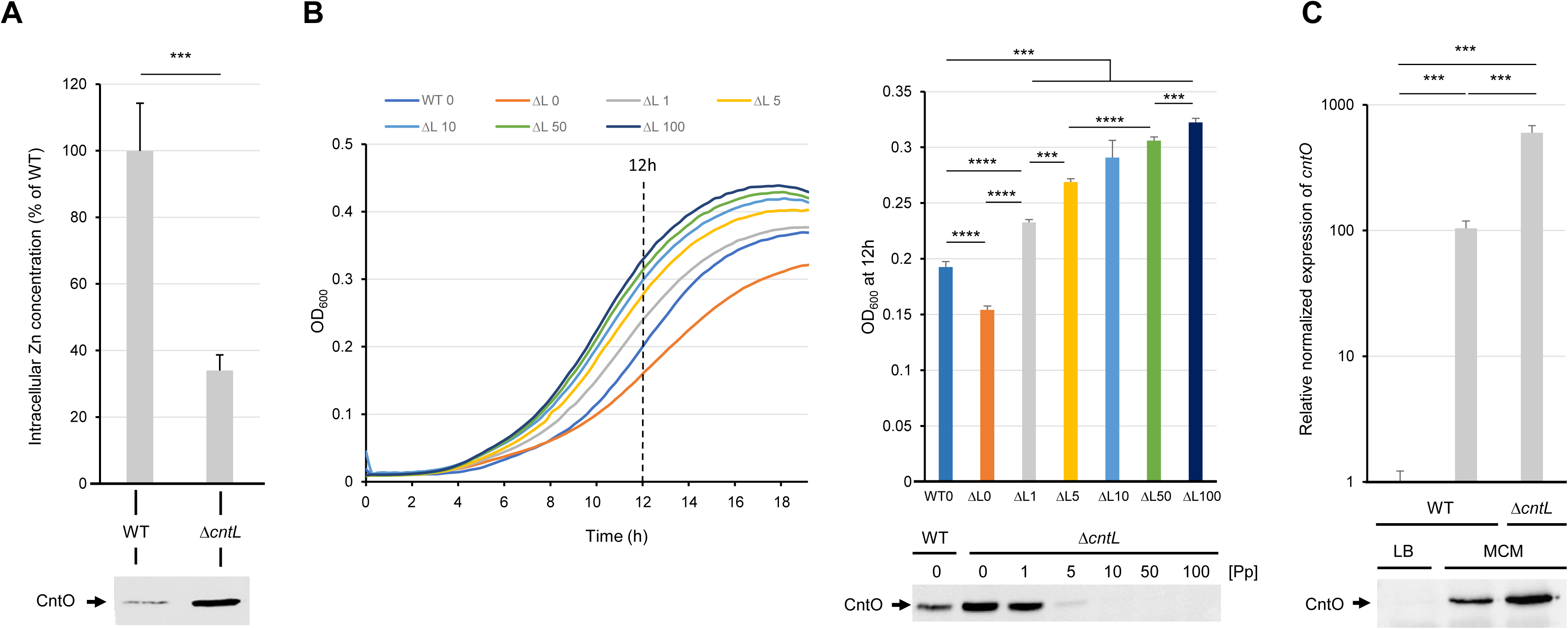
The pseudopaline pathway is induced and required for intracellular zinc homeostasis and optimal planktonic growth in MCM. (**A**) Impact of pseudopaline on intracellular zinc concentration and (**B**) cell growth of PA14 wild-type (WT) and *cntL* mutant (Δ*cntL* or Δ*L*) strains, supplemented or not with various concentration (in µM) of exogenous pseudopaline [Pp], grown in MCM. Complete growth curve and 12h point OD_600_ (dash line) are presented (**C**) qRT-PCR experiments revealing the direct correlation between *cntO* gene expression and CntO protein production in various growth conditions and genetic backgrounds. For all panels, CntO levels in the different protein samples were estimated by immunoblot experiments using CntO antibodies (lower panels). Error bars, mean ± standard deviation (sd) of at least three independent biological replicates. **** and *** correspond to p <.0001 and p <.001 respectively.

Additional experiments performed with different concentrations of externally added synthetic pseudopaline indicate that optimal WT growth is already achieved at 1 µM (Fig. 1B), revealing that in the metal limiting conditions encountered in MCM, the natural extracellular concentration of pseudopaline do not exceed 1 µM. Moreover, the observation that the addition of higher concentrations of pseudopaline leads to better growth than that observed in the WT also indicates that zinc recovered by pseudopaline under these conditions is the limiting factor for growth in MCM. Remarkably, the fact that growth of the mutant is already and completely restored upon addition of 1μM of exogenous pseudopaline also suggests that pseudopaline could be recycled, similar to the siderophore pyoverdine (Schalk et al., 2002). Thus, after recovery by the bacteria and release of its metal, pseudopaline could be secreted by the CntI and MexA/B-OprM transporters for a new zinc import cycle.

We might ask whether the alternative Znu zinc import pathway supports or at least contributes to the residual growth of the *cntL* mutant in MCM. The similar MCM generation times observed between the WT and the single *znuA* mutant on the one hand or the single *cntL* and the double *cntL*/*znuA* mutant on the other hand presented figure S1, indicate that this is not the case and that in MCM, another, poorly efficient zinc entry pathway that supports the low growth of *Pseudomonas* in absence of pseudopaline, remains to be discovered.

Additional transcriptional data, together with metal and protein level measurements, all presented figure 1, revealed a direct relationship between intracellular zinc concentration (Fig. 1A) and the levels CntO protein or the corresponding transcripts (Fig. 1C). Given the operon organization of the *cntOLMI* genes, encoding the OM receptor (*cntO*), the pseudopaline synthesis proteins (*cntL* and *cntM*) and the IM exporter (*cntI*) respectively (Lhospice et al., 2017), the direct relationship between intracellular zinc level and CntO protein production level can be extended to pseudopaline production level. Consequently, any intracellular zinc deficiency leads to an increase in pseudopaline production in order to secure proper zinc intracellular homeostasis establishing an additional interesting direct correlation between CntO production and pseudopaline essentiality.

Altogether, our data based on planktonic growth indicate that in MCM, zinc is scarce and only the secreted pseudopaline can import enough of this metal to sustain it at the intracellular levels needed to feed the zinc proteome and ensure optimal planktonic bacterial growth.

### Pseudopaline is required for proper biofilm formation in MCM

In order to evaluate the implication of pseudopaline in physiological functions relevant for *P. aeruginosa*’s fitness in its environment, including an infected host, we explored zinc acquisition in biofilm formation, the most common lifestyle reported during infections (Tuon et al., 2022). This multicellular organisation protects the bacterial community from external onslaughts such as phagocytosis or killing by antibiotics and allows for establishing chronic recalcitrant infections. We compared biofilm formation of WT and Δ*cntL* PA14 strains grown in MCM using several quantitative and qualitative approaches. We first examined biofilm formation in 24-wells polystyrene plates after 24h of static growth in MCM at 30°C. The *cntL* mutant, unable to produce pseudopaline, shows a significant 30% reduction of crystal-violet stained biofilm compared to the WT (Fig. 2A). Adding synthetic exogenous synthetic pseudopaline to the *cntL* mutant showed biofilm restoration, confirming that the role of pseudopaline in proper biofilm formation also comprises an extracellular stage. Cytoplasmic import of zinc by the exogenously added pseudopaline was confirmed by the repression of the *cnt* promoter, illustrated by the absence of CntO in the supplemented strain (Fig. 2A, bottom). The restoration of biofilm in the mutant by 100 µM pseudopaline to a level significantly higher than that observed in the WT also supports that, similar to the planktonic growth (Fig. 1B), pseudopaline-dependent zinc import is also the limiting factor for biofilm formation. These result shows that in *P. aeruginosa* biofilms grown under zinc-limited conditions, the zinc homeostasis required for its optimal formation is provided by pseudopaline.

**FIGURE 2.**
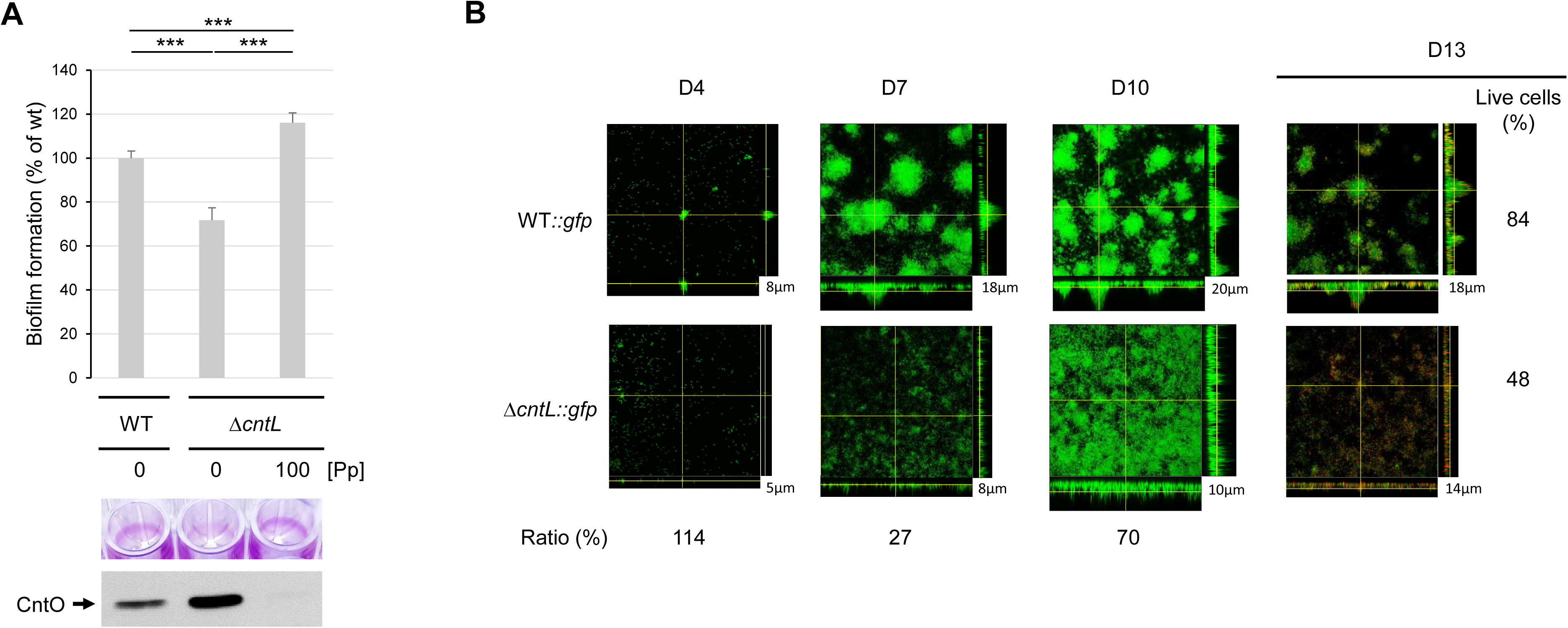
The pseudopaline pathway is induced and required for proper quantitative and qualitative biofilm formation in MCM. Impact of pseudopaline on biofilm formation of (**A**) PA14 wild-type (WT) and *cntL* mutant (Δ*cntL*) strains or (**B**), PA14 wild-type (WT::*gfp*) and *cntL* mutant (Δ*cntL*::*gfp*) Gfp-labelled strains, supplemented or not with exogenous pseudopaline (Pp) at 100µM and grown in MCM. (**A**) Biofilm formation was measured by quantifying bacterial mass adhering to the walls of a microtiter plate well as described in the M&M section. A representative illustration of crystal violet rings obtained with the different strains during the biofilm formation assay is presented in the middle panel. CntO productions in the different protein samples visualized by immunoblot experiments are presented in the lower panel. (**B**) Qualitative biofilm formation was monitored at days 4 (D4), 7 (D7) and 10 (D10) post-incubation. Stacked confocal scanning laser microscopy images of biofilms and corresponding extracted z images and their respective xy and xz planes as well as their thickness (in µm) are presented. The ratios (in %) indicate the average biomass of the *cntL* mutant (PA14Δ*cntL*::*gfp*) compared to the wild type (PA14wt::*gfp*) strains at the three different days of incubation. Antimicrobial tolerance of PA14wt::*gfp* and PA14Δ*cntL*::*gfp* strains 11 days aged biofilms, exposed to 20 μg/mL of tobramycin for 24 h and observed at day 13 (D13) after 24 h of flow. Cells with a compromised membrane that are dead or dying stain red with propidium iodide, whereas cells with an intact membrane stain green with GFP fluorescence. The live cells (in %) indicate the biomass of the live cells compared to the total live and dead cells obtained from two independent experiments. The experiment was repeated twice, independently, and the images presented correspond to the typical results observed in both experiments. Errors bar, mean ± standard deviation (sd) of at least three independent biological replicates. *** correspond to p <.001.

In order to assess the qualitative impact of the absence of pseudopaline on biofilm formation in MCM, we used confocal microscopy to follow its formation and organization in flow cell experiments using GFP-tagged PA14 strains. Biofilms formed by WT and Δ*cntL* PA14 strains were observed after 4, 7 and 10 days of incubation in a flow cell chamber. Biomass ratios and thickness measurements presented figure 2B show significant reductions in cell number and thickness of the biofilm in the absence of pseudopaline after 7 and 10 days of incubation. From a qualitative point of view, the *cntL* mutant formed biofilms with an impaired morphology compared to the parental WT strain, unable to form the classic pilar-and mushroom-shaped architecture (Fig. 2B). We further evaluate pseudopaline involvement in the antibiotic tolerance, a property described for *P. aeruginosa* biofilms (Costerton et al., 1999). D11 biofilms were treated for two days with 20 µg/mL of the aminoglycoside antibiotic tobramycin, generally used in inhalation in CF patients infected with chronic *P. aeruginosa* infection (Smith et al., 2022). The live (GFP-green) and dead (propidium iodine-red) detections applied at D13 highlight a significantly higher sensitivity to tobramycin for the *cntL* mutant compared to the WT (Fig. 2B right panels). This observation, not reported in planktonic growth where WT and *cntL* mutant strains were both sensitive from 3 µg/mL (Fig. S3) revealed that the absence of pseudopaline not only delays biofilm formation but also significantly impairs its structural properties and its ability to confer tolerance to an antibiotic. The negative effect of the absence of pseudopaline on the quantity and quality of the biofilm in metal-scarce conditions is most likely attributable to the impact of the reduced intracellular zinc level - reported by the higher amount of CntO detected (Fig. 2A lower panel) - on the proper function of bacterial zinc-dependant metalloproteins involved in this process (Marguerettaz et al., 2014).

The pseudopaline-dependent growth and biofilm phenotypes observed in this study (Figs. 1 & 2) highlight the broad importance of this metallophore in environments where zinc is scarce, such as the one created in MCM or reported during infections where nutritional immunity leads to serious metal deficiencies in general, and zinc in particular (Hood and Skaar, 2012; Kehl-Fie and Skaar, 2010; Vermilyea et al., 2021; Wakeman et al., 2016).

### Pseudopaline is required at different stages of macrophage infection

To further explore the role of the pseudopaline zinc uptake pathway at different steps of a host-cell infection process, we exploited RAW 264.7 macrophage-like murine cells (Felgner et al., 2020). We started by measuring phagocytic uptake through the gentamycin-protection assay and observed that the PA14Δ*cntL* mutant strain pre-cultured in MCM is significantly less phagocyted than the parental PA14 WT strain, a phenotype mostly recovered in the PA14Δ*cntL::cntL* complemented strain (Fig. 3A). We hypothesized that this reduced phagocytic uptake could be due to a reduced adherence of the *cntL* mutant. To address this hypothesis, we blocked phagocytosis by adding 10 µM cytochalasin D to the culture medium before infection, and determined adherent bacterial cells to the RAW 264.7 membrane. We observed that the *cntL* mutant indeed showed a decreased adherence to the macrophages (Fig. 3B), thus explaining, at least in part, the reduced phagocytic uptake.

**FIGURE 3.**
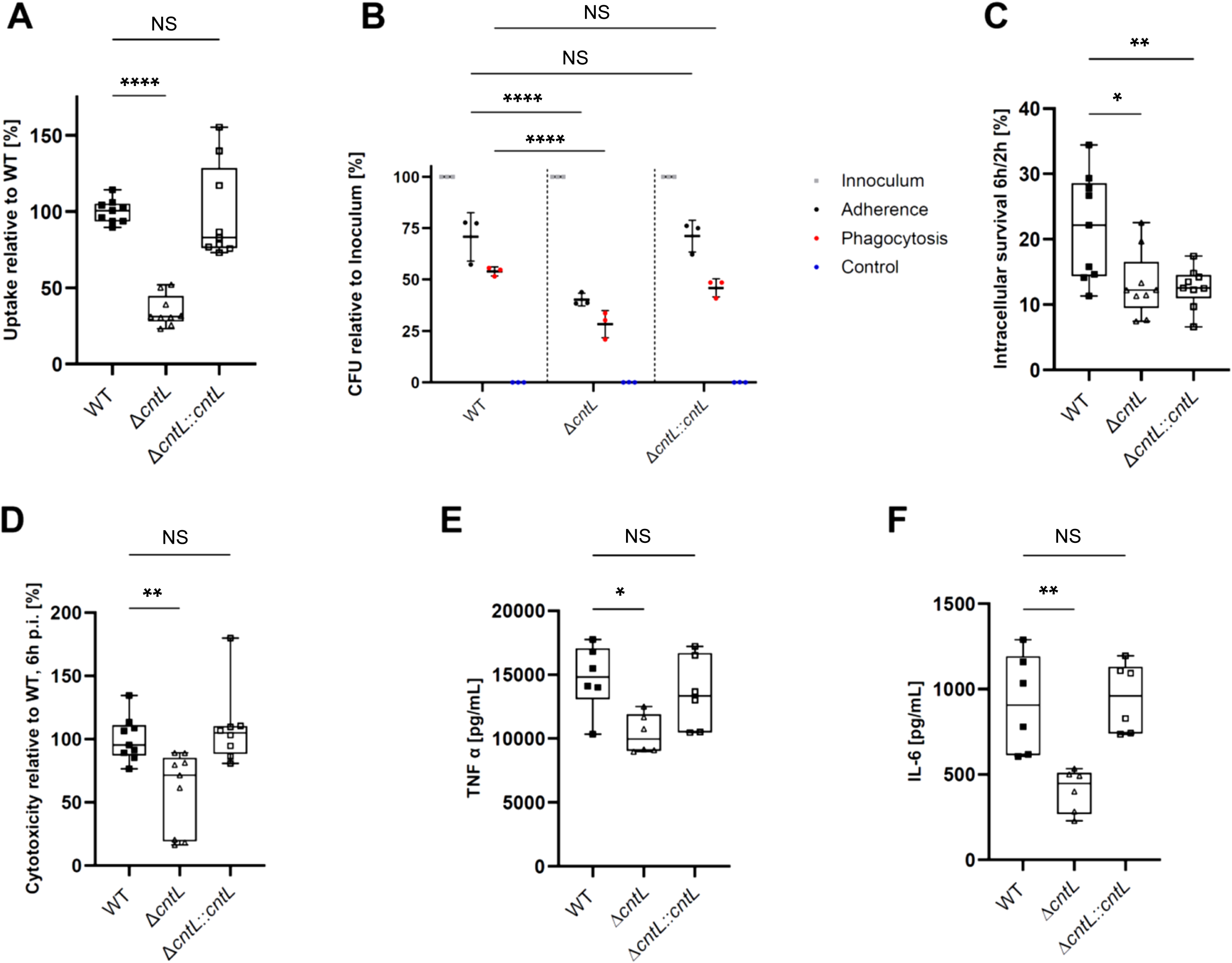
The pseudopaline pathway is required at different stages of macrophage infection by *P. aeruginosa*. Strains PA14 WT (WT), *cntL* mutant (Δ*cntL*) and *cntL cis-*complemented mutant (Δ*cntL*::*cntL*) precultured in MCM were compared in different macrophage infection essays. (**A**) Phagocytic uptake into RAW 264.7 cells. The results of three independent experiments with three biological replicates are presented. **** correspond to p<0.0001 (One-way ANOVA, Tukey Post-hoc test). (**B**) Adherence measurement to RAW 264.7 cells, following treatment of cells with cytochalasin D to stop phagocytosis. The inoculum (grey) were set as 100% for each strain and the uptake without cytochalasin D (corresponding to phagocytosis) was also run as internal control (red), as well as a negative control with cytochalasin and gentamicin in which no CFU were recovered (blue). Mean ± standard deviation of three biological replicates are shown. Phagocytic uptake **** correspond to p<0.0001 (Two-way ANOVA, Dunnett Post-hoc test). (**C**) Bacterial survival in RAW 264.7 macrophages. Macrophages were infected by *P. aeruginosa* strains and treated with gentamicin 1 hpi. Ratio of CFU recovered 6 hpi were normalized to CFU recovered 2 hpi, and expressed as percentage. The results of three independent experiments with three biological replicates are depicted. * and ** correspond to p<0.0332 and p<0.0021 respectively (One-way ANOVA, Tukey Post-hoc test). (**D**) Cytotoxicity on RAW 264.7 macrophages. Cells were infected at MOI of 1 after culture in MCM and treated with gentamicin 1 hpi. Supernatant was sampled 6 hpi, and lactate dehydrogenase (LDH) activity measured by a colorimetric assay. Values are normalized to LDH activity upon infection with PA14 WT strain, expressed as percentage. The results of three independent experiments with three biological replicates are depicted. ** correspond to p<0.0021 (One-way ANOVA, Tukey Post-hoc test). Quantification of Tumor necrosis factor α (TNF-α) (**E**) and interleukin 6 (IL-6) (**F**) produced 6 hpi by RAW 264.7 macrophages infected by *P. aeruginosa* strains at MOI 1 after growth in MCM and treated with gentamycin 1 hpi. The results of two independent experiments with three biological replicates are depicted. * and ** correspond to p<0.0332 and p<0.0021 respectively (One-way ANOVA, Tukey Post-hoc test).

Once phagocyted, bacterial cells undergo numerous stresses imposed by the macrophage for its clearance. We evaluated pseudopaline involvement in bacterial survival by measuring the number of colony-forming-units (CFU) recovered 6 hours post-infection (hpi). To account for the decreased phagocytosis of the *cntL* mutant compared to the WT, we normalized the survival to the number of CFU phagocyted 2 hpi. Under these phagocytosis-independent conditions, the *cntL* mutant shows a significant reduced survival in the macrophages than the WT (Fig. 3C). This phenotype is accompanied by both a reduced cytotoxicity (Fig. 3D) and a decreased cytokine production of the *cntL* mutant compared to the WT, as determined by measuring levels of TNF-α (Fig. 3E) and IL6 (Fig. 3F) produced by the RAW 264.7 cells. In these macrophage infection experiments, all but one phenotypes due to *cntL* deletion are properly complemented by the *cntL* allele, cloned at the *att* site on the chromosome. The lack of complementation of the intracellular survival phenotype shown in Figure 3C may be due to the significant reduction of pseudopaline in the supernatant of the complemented strain, previously reported by Lhospice et al. (Lhospice et al., 2017) and confirmed in the present study by the more than 6-fold reduction in *cntL* transcription in the complemented strain (Figure S4). The reduced amount of extracellular pseudopaline in the complemented strain appears to be only relevant for the intracellular survival phenotype for which an optimal level of extracellular pseudopaline would be mandatory to counteract - by metal captation in the extracellular medium - bacterial intoxication by metals other than zinc in the macrophage vacuole (Palmer and Skaar, 2016).

These results demonstrate the critical role of pseudopaline at different stages of infection in macrophages, supporting the diversity of the pool of zinc-dependant metalloproteins involved in this process whose metal supply depends on pseudopaline. They also confirm that the nutritional zinc deficiency imposed on the pathogen by the macrophage, requires the induction of the pseudopaline pathway, which therefore proves to be mandatory for the expression of its full pathogenic power throughout the macrophagic infection.

### Pseudopaline plays a critical role during *P. aeruginosa* pulmonary mouse infection

To further extend our cell culture-based findings in an *in vivo* animal model with a fully functional immune system model, we used a well-established mouse pulmonary infection model (Bragonzi, 2010). Mice were infected by pulmonary instillation of MCM-grown WT and Δ*cnt* PA14 *P. aeruginosa*. Several markers including body weight, clinical score and survival were recorded during the infection. While all mice infected with the WT suffered from severe conditions characterized by continuous body-weight loss (Fig. 4A), increased clinical score (Fig. 4B) and > 50% of death 4 days post-infection (Fig. 4C), the mice infected with the *cnt* mutant exhibited significantly milder evidence of susceptibility, including regain of body-weight loss 2 days post-infection (Fig. 4A), less sever clinical symptoms (Fig. 4B) and almost full survival after 7 days post-infection (Fig. 4C). To gain insight into the mechanisms that contributed to the lower virulence associated with the *cnt* mutant, we analysed the host’s antibacterial responses 48 hpi. Mice infected with the WT showed significantly higher bacterial load in broncho-alveolar fluid (BAL) and entire lungs as compared to *cnt* mutant infected animals (Figure 4D). Given that pneumonia is intimately associated with lung inflammation, we evaluated the production of IL6, a prototypal pro-inflammatory cytokine, in the BAL and in the lungs (Fig. 4E). As observed with macrophage infections, the present analysis revealed that IL6 was significantly reduced in the animals infected by the *cnt* mutant as compared to the WT. Altogether, these data confirm attenuation of virulence of the pseudopaline-deficient *P. aeruginosa* strain in a mouse model of lung infection.

**FIGURE 4.**
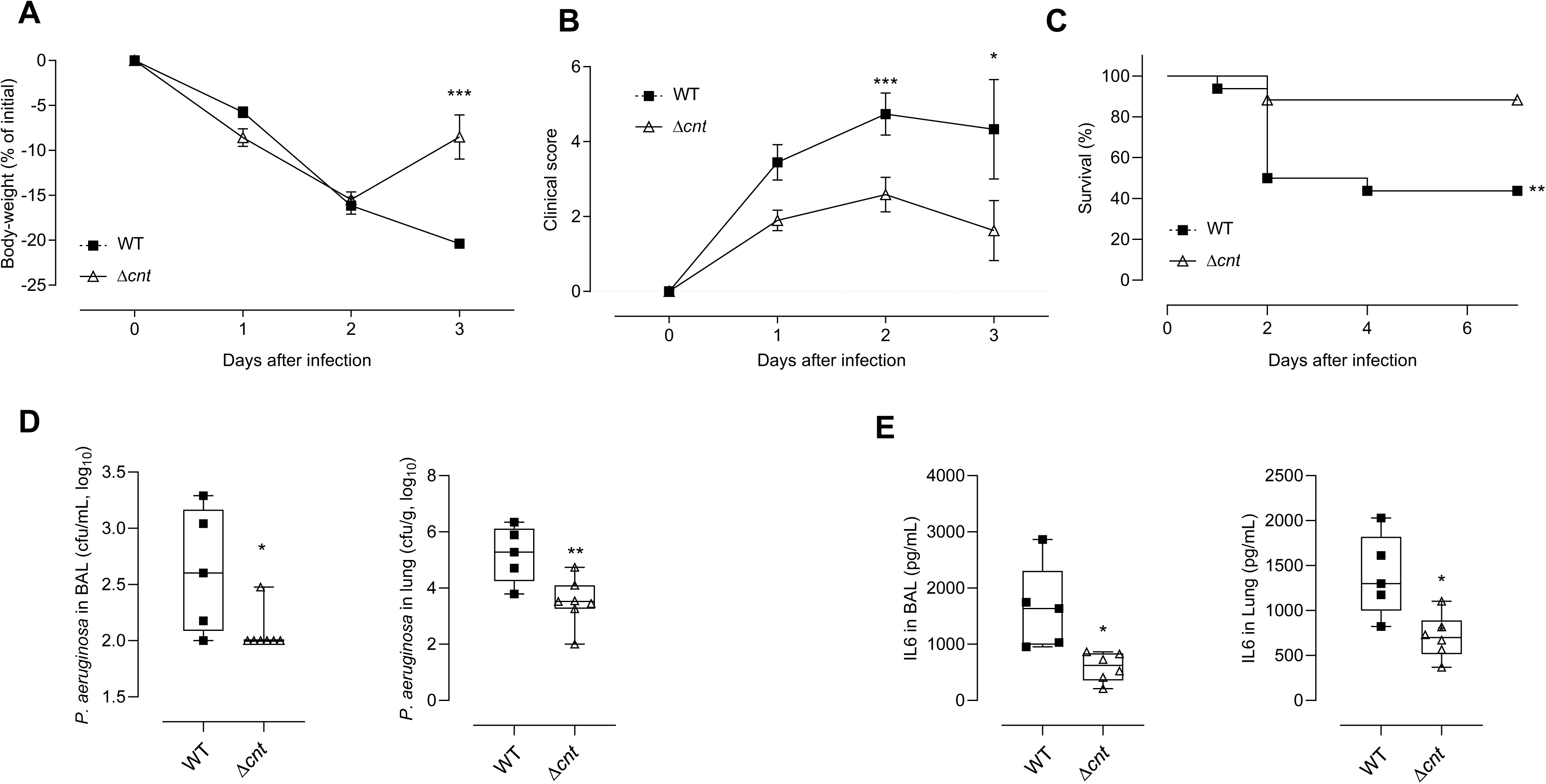
The pseudopaline pathway plays an important role during *P. aeruginosa* pulmonary mouse infection. Mice were infected intratracheally with PA14 wild-type (WT) and *cnt* mutant (Δ*cnt*) strains precultured in MCM. Animal weight **(A)**, clinical score **(B)** and survival **(C)** were recorded throughout the experiment as described in the M&M section. (**D**) In a separate experiment, mice were sacrificed at 2 days post-infection and the number of PA14 in the bronchoalveolar lavage fluid (BAL) and lungs was determined. **(E)** The amount of interleukin 6 (IL-6) was measured in the bronchoalveolar lavage and lungs. Each result corresponds to at least 3 independent experiments (n=17/group) for A-C and 1 experiment (n=5-7/group) of D and E.

The phenotype of pseudopaline-deficient *P. aeruginosa* in the mouse infection model indicates an important role for the pseudopaline Cnt pathway in pathogenesis of this organism. This is consistent with the pseudopaline-dependent survival of *P. aeruginosa* in airway mucus secretions reported by Gi and colleagues (Gi et al., 2015). The numerous mice or human transcriptome data showing a systematic up-regulation of the *cnt* genes under similar pulmonary infection conditions therefore extend the role played by the pseudopaline to pulmonary infections in general (Cornforth et al., 2018; Mastropasqua et al., 2018; Turner et al., 2015). The pseudopaline pathway is therefore important for *P. aeruginosa* survival in a mammalian host when it must cope with severe metal deficiencies such as those encountered in pulmonary infections and possibly due to sequestration by calprotectin (Neff et al., 2024).

### Conclusion

Results presented in this study revealed that under metal scarce and chelating conditions such as the one encountered during pulmonary infections, among the two zinc import pathways described in *P. aeruginosa,* the Cnt pseudopaline zinc uptake pathway proves to be the last resort and unique pathway allowing sufficient zinc uptake into the bacterium to preserve the proper intracellular zinc concentration necessary to maintain the pool of zinc metalloproteins fully functional. We are reporting in this study numerous phenotypes affected by zinc privation due to the absence of pseudopaline, that can be explained by the presence of a large number of zinc metalloproteins, previously shown to be important for many cellular functions including transcription, translation, motility or protease activity (Mikhaylina et al., 2018; Palmer and Skaar, 2016; Wang et al., 2021). It is noteworthy that the pseudopaline pathway, which is remarkably conserved in all strains of *Pseudomonas aeruginosa*, including clinical isolates, is absent in other closely related *Pseudomonas* species that colonize soil, water and plants, such as *P. fluorescens*, *P. putida*, *P. stutzeri* and *P. syringae*. Therefore, it can be assumed that pseudopaline is particularly important for promoting zinc recruitment in the context of animal hosts. The identification of the essential and multifactorial role of pseudopaline in *Pseudomonas* infections makes this pathway an interesting alternative target for the development of new antibacterials against *P. aeruginosa* infections.

## MATERIAL & METHODS

### Bacterial strains and preculture or planktonic growth conditions

Bacterial strains used in this study are listed in Table 1. All *P. aeruginosa* strains used in this study were derivatives of the parental PA14 strain. In macrophage infection experiments, the absence of *ctnL* in the PA14Δ*cntL* strain was complement by *cis* chromosomal expression of the *cntL* gene under the *cnt* operon promotor, inserted at the *att* site of the *P. aeruginosa* chromosome (Lhospice et al., 2017). Unless otherwise specified, precultures and planktonic growths of all *P. aeruginosa* strains were performed in minimal chelated medium MCM - minimal succinate (MS) media (per liter: 30g K_2_HPO_4_, 15g KH_2_PO_4_, 5g (NH_4_)_2_SO_4_, 1g MgSO_4_, 20g Succinic acid and 15.5g NaOH, pH 7.0, MS) supplemented with 100µM EDTA - at 37 °C with horizontal shaking in polycarbonate sterile single-used Erlenmeyer flasks or 96 well plates in order to avoid any metal contamination and monitored by OD_600_ measurements.

**Table 1:** Bacterial strains used in this study.

| *P. aeruginosa* Strains | Descriptions | References |
| --- | --- | --- |
| PA14 WT | Wild type strain | (Liberati et al., 2006) |
| PA14 WT::*gfp* | Wild type PA14 strain harboring the *gfp* gene under constitutive promotor at *att* site | (This study) |
| PA14$\Delta cntL$ | *cntL* deletion mutant | (Lhospice et al., 2017) |
| PA14$\Delta cntL::gfp$ | PA14$\Delta cntL$ strain harboring the *gfp* gene under constitutive promotor at *att* site | (This study) |
| PA14$\Delta cntL::cntL$ | PA14$\Delta cntL$ harboring a wild type copy of *cntL* gene at *att* site. | (Lhospice et al., 2017) |
| PA14 $\Delta znuA$ | *znuA* deletion mutant | (This study) |
| PA14 $\Delta cntL \Delta znuA$ | *znuA cntL* double deletion mutant | (This study) |
| PA14$\Delta cnt$ | PA14 strain deleted for the complete *cntOLMI* operon. | (Gomez et al., 2021) |

### *Construction of* znuA *and* znuA-cntL *deletion mutant strains of* P. *aeruginosa*

Two DNA fragments corresponding to upstream and downstream regions of *znuA* gene were amplified from PA14 chromosomal DNA with PCR primers Mut-*znuA*-1/Mut-*znuA*-2 and Mut-*znuA*-3/Mut-*znuA*- 4 (Supplementary Table 2). Upstream and downstream regions were ligated by overlapping PCR and cloned into linearized pKNG101(Kaniga et al., 1991) by the SLIC method. The resulting constructs were transformed into *E. coli* CC118λpir and introduced into *P. aeruginosa* PA14 by conjugation. The strains in which the chromosomal integration event occurred were selected on *Pseudomonas* isolation agar plates supplemented with gentamycin. Excision of the plasmid, resulting in the deletion of the chromosomal target gene was performed after selection on Luria-Bertani (LB) plates containing 6% sucrose. Sucrose resistant and streptomycin sensitive clones were confirmed to be deleted for *znuA* gene by PCR analysis using Mut-*znuA*-5/Mut-*znuA*-6. The Δ*znuA* Δ*cntL* double mutant strain was constructed by knocking-out *znuA* in the *cntL* mutant strain.

### Pseudopaline supplementation experiments

MCM cultures of the pseudopaline deficient strain (PA14Δ*cntL*) used in planktonic and biofilm growths were supplemented with various externally added concentrations of chemical S-pseudopaline synthetized as described in (Zhang et al., 2019) and resuspended in ultrapure water at 100 mM stock solution.

### RNA Preparation and Reverse Transcription

RNAs were prepared from 25 mL culture of PA14 grown in LB or MCM after 10h shaking at 37°C. The cells were harvested by centrifugation and frozen at-80°C. Total RNAs were isolated from the pellet using RNeasy mini Kit (Qiagen) according to the manufacturer’s instructions; an extra TURBO DNAse (Invitrogen) digestion step was done to eliminate the contaminating DNA. The RNA quality was assessed by the TapeStation 4200 system (Agilent). RNA was quantified spectrophotometrically at 260 nm (NanoDrop 1000; Thermo Fisher Scientific). For cDNA synthesis, 1 µg total RNA and 0.5 μg random primers (Promega) were used with the GoScript™ Reverse transcriptase (Promega) according to the manufacturer’s instruction.

### Quantitative real-time retro transcriptional PCR (qRT-PCR)

qRT-PCR analyses were performed on a CFX96 Real-Time System (Bio-Rad, France). The reaction volume was 15 μL and the final concentration of each primer was 0.5 μM. The cycling parameters of the qRT-PCR were 98°C for 2 min, followed by 45 cycles of 98°C for 5 s, 60°C for 10 s. A final melting curve from 65°C to 95°C was added to check the specificity of the amplification. To determine the amplification kinetics of each product, the fluorescence derived from the incorporation of EvaGreen into the double-stranded PCR products was measured at the end of each cycle using the SsoFast EvaGreen Supermix 2X Kit (Bio-Rad). The results were analyzed using Bio-Rad CFX Manager software, version 3.1 (Bio-Rad). The 16S RNA and *uvrD* genes were used as a reference for normalizations. For each point a technical triplicate was performed. The amplification efficiencies for each primer pairs were comprised between 80 and 100%. All of the primer pairs used for qRT-PCR are shown in the table 2.

**Table 2:**
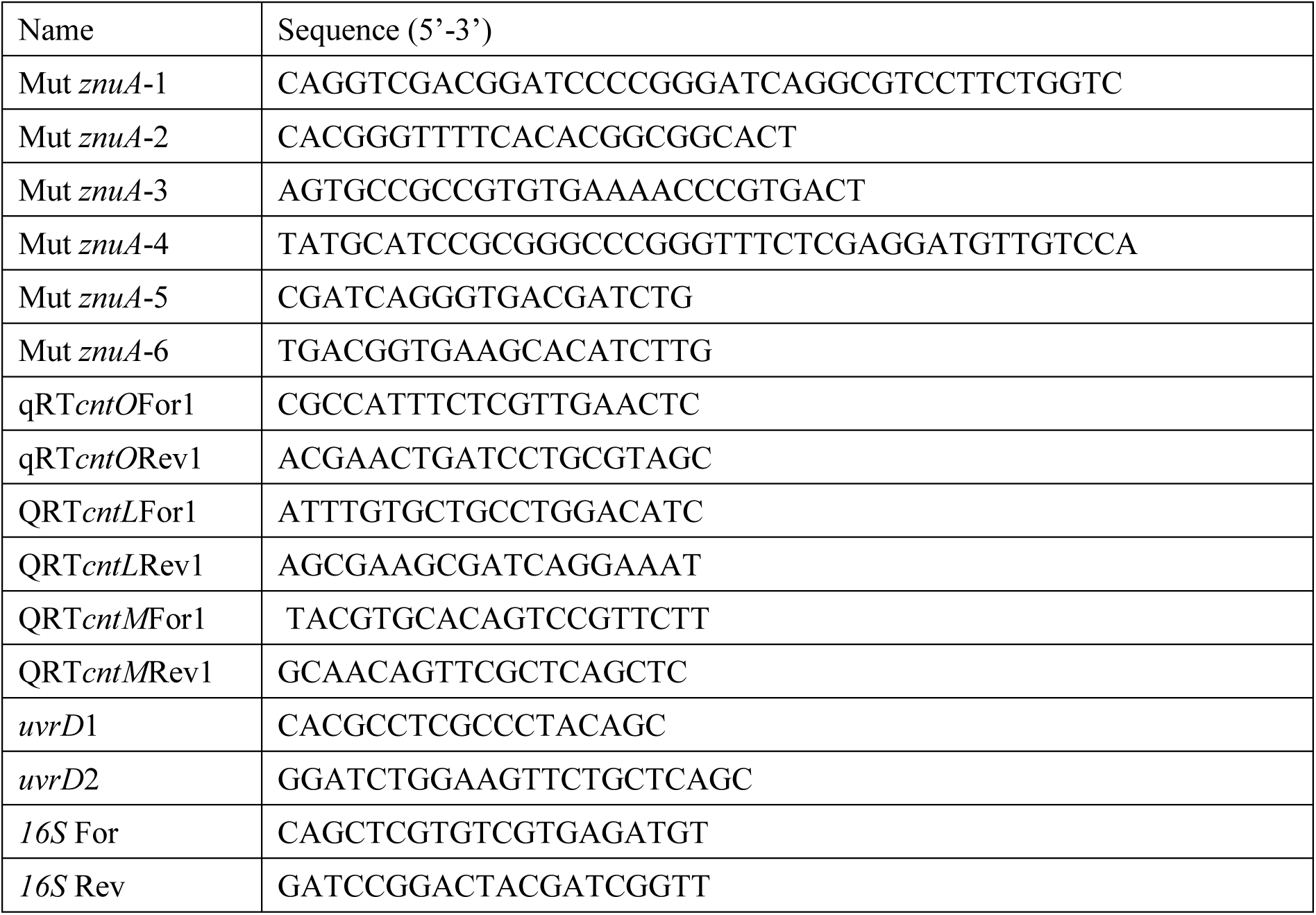
Primers used in this study.

### Biofilm formation

Biofilm formation for quantitative evaluation was measured in clear, flat bottom, 24-well plates (Corning^TM^ Costar 3526). Each overnight culture were inoculated at OD_600_ of 0.2 in 1 mL fresh MCM and incubated at 30°C for 24 hours. Following incubation, planktonic bacteria and media were rinsed away with non-sterile deionized water. The wells were filled with 0,1% crystal violet solution and after 15min at room temperature, they were washed three times with deionized water. Any crystal violet staining on the bottom of the well were cleaned, and the plate was let to air-dry overnight at room temperature. Crystal violet rings were then solubilized in 30% glacial acetic acid. Adherent biofilm was quantified by measuring optical density of all samples at 550 nm and normalized to mean absorbance from wells containing WT strain. Images used for biofilm illustration are representative of three independent experiments.

Time-lapse qualitative biofilm formation was performed in flow chambers as previously described (Giraud et al., 2011). Each experiment was repeated four times. The *P. aeruginosa* PA14 WT*::gfp* and PA14Δ*cntL::gfp* were tagged with green fluorescent protein (GFP), following the procedure describe in (Koch et al., 2001) except that conjugation instead of electroporation was used to transfer plasmids into *P. aeruginosa*. Bacteria from MS overnight precultures were inoculated at OD_600_ of 0.1 in MCM in flow chambers with individual channels (dimensions of 1 × 4 × 40 mm^3^) and incubated for 3h at 30°C before flow start. At day 11 (D11), tobramycin at 20µg/mL was added and incubated for 24 h with stopped flow. At day 12, the flow was turned on for 30 min, before addition of propidium iodide and incubation for 30 min with stopped flow. The flow was started again for 24 h before observation at day 13. Observation was performed after 4, 7, 10 and 13 days with an Olympus FV-1000 microscope equipped with detectors and a filter set for monitoring GFP and propidium iodide. The average biofilm thickness and biomass were calculated using Fiji (Schindelin et al., 2012).

### Western blot analysis

For analysis of CntO production, protein samples from bacteria corresponding to 1 OD unit were resuspended in 50µl 1X SDS-PAGE loading Laemmli buffer containing β-mercaptoethanol, heated at 95 °C for 10 min and separated on 12% acrylamide SDS-PAGE. After separation, the proteins were transferred onto nitrocellulose membranes for western blot analysis. Transfers were performed with semi-dry technic using Power Blotter Station (Invitrogen). The membranes were washed briefly in Tris-buffered saline, 0.10 % Tween 20 (TBS-T) and then blocked in TBS-T + 10 % milk for 1 hour at RT. The membranes were incubated for 1 h at RT in TBS-T + 10 % milk containing the primary (rabbit) antibody directed against CntO (laboratory collection) at 1/1,000 dilution. This was followed by 4 washes in TBS-T before an incubation of 1 h at RT in TBS-T + 10 % milk containing the secondary antibody anti-mouse-HRP conjugated (1/5,000). Finally, after five more washes with TBS-T, the peroxidase reaction was revealed by chemiluminescence (SuperSignal™ West Dura Extended Duration Substrate, Thermo Scientific) and scanned with ImageQuant LAS 4,000.

### Macrophage infections

#### Phagocytic uptake and survival

RAW264.7 (ATCC number: TIB-71TM) cells were routinely grown in DMEM (4.5 g/l Glucose) and supplemented with 10% FCS, 20 mM HEPES and 1% BSA at 37°C, 5% CO_2_. All cell cultures used in this study were tested and confirmed as mycoplasma-free. Invasion assays were performed as described previously (Felgner et al., 2020). Briefly, 5×10^5^ RAW 264.7 cells were seeded in DMEM in a 24 well plate for phagocytic uptake or intracellular replication measurements. Before the start of the experiments, the supernatant of each well was discarded and replaced with 400 μL of fresh DMEM without phenol red. Eukaryotic cells were infected at a MOI (multiplicity of infection) of 1 with selected *P. aeruginosa* strains grown 10 h in MCM, washed with PBS, and adjusted to a final volume of 100 μL in fresh DMEM. Number of CFU in the inoculum dose is verified by serial plating on LB plates. Immediately after start of the infection, a centrifugation step at 1200 x *g* for 5 min was used to ensure cell-to-cell contact. To kill extracellular bacteria, one-hour post-infection (hpi), 100 μL of DMEM containing gentamicin was added to the final concentration of 50 μg/mL. At 2hpi, cells were washed with PBS and lysed with PBS containing 0.5% (v/v) TritonX-100. CFUs were determined by plating of serial dilutions and compared to the parental strains. These CFU represent the number of cells phagocyted, and normalized to the inoculum. A replicate plate was used for survival assay, using the same inoculum and cells batch. After 6h, cells were washed with PBS and lysed with PBS containing 0.5% (v/v) TritonX-100. CFUs were determined by plating of serial dilutions and compared to the parental strains. Intracellular survival is evaluated by the ratio of CFU recovered after 6h, and normalized to the number of cells phagocyted after 2 h.

#### Adherence assay

Adhesion assay was performed as described before (Felgner et al., 2020). Briefly, RAW234.7 cells were seeded in DMEM as detailed in the phagocytic uptake and survival assays, and infected by *P. aeruginosa* strain grown in MCM for 10 h. 10 µM of Cytochalasin D was added to cell cultures 2 h prior to infection to block phagocytosis. Directly after infection, the bacteria were centrifuged at 1200 x g for 5 min to ensure cell-to-cell contact. 2 hpi, the supernatant was removed. The cells were then gently washed twice with PBS and lysed using PBS containing 1% (v/v) TritonX-100. Serial dilutions were used to determine the adherent bacteria by plating and counting CFUs.

#### Cytotoxicity

50 μL of supernatant of infected RAW264.7 cells were collected 6 hpi and LDH levels were measured using the CytoTox 96 Non-radioactive Cytotoxicity Assay (Promega) according to the manufacture’s protocol. PBS serves as negative control and lysis solution of the manufacturer was used as killing control. Values are expressed as % of the value obtained with the PA14 WT strain infection.

#### Measure of cytokine production

Macrophage supernatants were sampled 6 hpi. The TNF-α and IL-6 ELISA Max Standard Kit (Biolegend) were used to determine the TNF-α and IL-6 levels according to the manufacturer’s manual. Three different biological replicates were analysed in technical triplicates, and a PBS-treated group served as negative control.

### Pulmonary mouse infections

#### Mice

Adult male C57BL/6jrj (B6) mice (6-8 weeks old) were obtained from Janvier (Le Genest Saint-Isle, France). All mice were housed under specific-pathogen-free conditions at the “PST Animaleries” animal facility (Université de Tours, France) and had access to food and water *ad libitum*. All experiments complied with the French government’s ethical and animal experiment regulations (APAFIS#7608- 2016111515206630).

#### P. aeruginosa infections

MCM precultures of WT and Δ*cnt* PA14 *P. aeruginosa* strains were used in this study. Mice were anesthetized with isoflurane 4% and an operating otoscope fit with intubation specula was introduced to both maintain tongue retraction and visualize the glottis. A fiber optic wire threaded through a 20G catheter and connected to torch stylet (Harvard Apparatus, France) was inserted into the mouse trachea. Correct intubation was confirmed using lung inflation bulb test and 40µL of the bacterial inoculum (2- 8×10^6^ cfu) was applied using an ultrafine pipette tip. Inoculum size for infections were confirmed after each experiment by counting colony-forming unit (cfu) on *Pseudomonas* isolation agar plates. Mortality, body weight and clinical score of animals were monitored daily. Clinical score was calculated by recording the following symptoms: ruffled fur, hunched posture, motor impairment, breathing impairment, closed eyes, purulent eyes and allocating 1 point to each. In all experiments, moribund animals - animals with a weight loss of more than 20% or clinical score of 7 - were sacrificed for ethical reasons and considered as dead due to the infection.

#### Broncho-alveolar lavage, organ sampling and bacterial load assay

At day 2 after infection, mice were sacrificed with intra-peritoneal injection of pentobarbital. Broncho-alveolar lavage fluid (BAL) was collected after *P. aeruginosa* infection by introducing a catheter into the trachea under deep pentobarbital anaesthesia and washing sequentially the lung with 2 x 1mL of 1X PBS at room temperature. The lavage fluid was centrifuged at 400 g for 10 min at 4°C and the supernatant was stored at-20°C for ELISA analysis. Lungs were perfused with 10mL of PBS1X and harvested in GentleMACS M tubes (Miltenyi Biotec, Germany) containing 2 mL of 1X PBS for microbiology assay. Bacterial load in lung homogenates or BAL (before centrifugation) was determined by plating tenfold serial dilutions on *Pseudomonas* Isolation Agar (PIA) plates were incubated at 37°C in a 5% CO_2_ atmosphere, and the CFUs were counted after 24 hours.

#### Measure of cytokine production

Concentration of IL-6 in BAL supernatants and lung homogenates were measured using specific ELISA Max Standard Kit (Biolegend) according to the manufacturer’s manual.

### Statistics

Statistics were determined using the Student’s t-test function of excel using a bilateral model and assuming equal variance when distribution of samples had normal distribution (Fig. 1). For nonparametric data (Figs 2, 3 and 4) statistical significance was calculated using the Wilcoxon rank sum test. Statistical evaluation of differences between the experimental groups was determined by using two-way analysis of variance (ANOVA) followed by a Sidak’s post-test (which allows comparison of all pairs of groups) (Fig. 4). Log-rank test was used for survival analysis (Fig. 4).

## ACKNOWLEDGEMENTS

We gratefully to Pr. Steve Lory for careful reading of the Manuscript. We also thank the whole RV group for constant support in the project. This work was supported by the grants: RF20210502871 from “Vaincre la Mucoviscidose” and “Gregory Lemarchal” associations allocated to RV, NHV and TS; 2019-00131897 NOVANTINH from the “Région Centre-Val-de-Loire” allocated to NHV, TS and RV and ANR-20-CE11-0018 “Pseudo-Traffic” allocated to RV and PA. S.H. received funding from the DFG in the DFG SPP 1879 program, under Germany’s Excellence Strategy – EXC 2155 “RESIST”— Project ID 390874280, from the Lower Saxony Ministry for Science and Culture (BacData ZN3428), and from the Novo Nordisk Foundation (NNF 18OC0033946).

## AUTHOR CONTRIBUTIONS

Conceive the experiments: NOG, TS, LB, MB, RV. Perform, analyse and interpret the experiments: NOG, TS, LB, GB, DLP, MT, MB, YD, LO, PA, MB, CB, RV. Write the paper: NOG, RV. Review the paper: TS, LB, GB, MT, PA, LO, XL, YB, SH, NHV. Secure funding’s: NHV, TS, SH, PA and RV.

The authors declare no conflict of interest

## FIGURE LEGENDS

**FIGURE S1**

**The Znu pathway does not support *Pseudomonas* growth in MCM.** Generation time of PA14 wild-type (WT), *cntL* mutant (Δ*cntL*), *znuA* mutant (*ΔcntznuA*) or *cntL/znuA* double mutant (Δ*cntL*Δ*znuA*) strains, grown in MCM. Error bars, mean ± standard deviation (sd) of at least three independent biological replicates. *** correspond to p <.001; NS for not statistically significant.

**FIGURE S2**

***cntL* deletion does not affect *cntM* expression in MCM.** qRT-PCR experiments revealing the similar level of expression of *cntM* in PA14 wild-type (WT) and *cntL* deletion mutant (Δ*cntL*) grown in MCM thus excluding any polar effect of the *cntL* deletion. Error bars, mean ± standard deviation (sd) of at least three independent replicates.

**FIGURE S3**

**Tobramycin sensitivity assay of PA14 WT and *cntL* mutant strains grown in MCM.** Cell growth in MCM of PA14 wild-type (WT) and *cntL* mutant (Δ*cntL*) strains grown in presence of increased concentration of tobramycin.

**FIGURE S4**

***cntL* expression is significantly reduced in the PA14 Δ*cntL::cntL* complemented strain.** qRT-PCR experiments revealing the more than 6 time reduction of *cntL* expression in PA14 Δ*cntL::cntL* strain compared to the PA14 Δ*cntL* strain under similar MCM growth conditions. Error bars, mean ± standard deviation (sd) of at least three independent replicates.

## REFERENCES

Andreini, C., Banci, L., Bertini, I., and Rosato, A. (2006). Zinc through the three domains of life. J Proteome Res 5, 3173–3178.

Andreini, C., Bertini, I., and Rosato, A. (2009). Metalloproteomes: a bioinformatic approach. Acc Chem Res 42, 1471–1479.

Bertini, I., Sigel, A., and Sigel, H. (2001). Handbook on Metalloproteins (New York).

Bielecki, P., Puchalka, J., Wos-Oxley, M.L., Loessner, H., Glik, J., Kawecki, M., Nowak, M., Tummler, B., Weiss, S., and dos Santos, V.A. (2011). In-vivo expression profiling of Pseudomonas aeruginosa infections reveals niche-specific and strain-independent transcriptional programs. PLoS One 6, e24235.

Bragonzi, A. (2010). Murine models of acute and chronic lung infection with cystic fibrosis pathogens. Int J Med Microbiol 300, 584–593.

Cornforth, D.M., Dees, J.L., Ibberson, C.B., Huse, H.K., Mathiesen, I.H., Kirketerp-Moller, K., Wolcott, R.D., Rumbaugh, K.P., Bjarnsholt, T., and Whiteley, M. (2018). Pseudomonas aeruginosa transcriptome during human infection. Proc Natl Acad Sci U S A 115, E5125–E5134.

Costerton, J.W., Stewart, P.S., and Greenberg, E.P. (1999). Bacterial biofilms: a common cause of persistent infections. Science 284, 1318–1322.

Cunrath, O., Geoffroy, V.A., and Schalk, I.J. (2016). Metallome of Pseudomonas aeruginosa: a role for siderophores. Environ Microbiol 18, 3258–3267.

D’Orazio, M., Mastropasqua, M.C., Cerasi, M., Pacello, F., Consalvo, A., Chirullo, B., Mortensen, B., Skaar, E.P., Ciavardelli, D., Pasquali, P., et al. (2015). The capability of Pseudomonas aeruginosa to recruit zinc under conditions of limited metal availability is affected by inactivation of the ZnuABC transporter. Metallomics 7, 1023–1035.

Ellison, M.L., Farrow, J.M., 3rd, Parrish, W., Danell, A.S., and Pesci, E.C. (2013). The transcriptional regulator Np20 is the zinc uptake regulator in Pseudomonas aeruginosa. PLoS One 8, e75389.

Felgner, S., Preusse, M., Beutling, U., Stahnke, S., Pawar, V., Rohde, M., Bronstrup, M., Stradal, T., and Haussler, S. (2020). Host-induced spermidine production in motile Pseudomonas aeruginosa triggers phagocytic uptake. Elife 9.

Gi, M., Lee, K.M., Kim, S.C., Yoon, J.H., Yoon, S.S., and Choi, J.Y. (2015). A novel siderophore system is essential for the growth of Pseudomonas aeruginosa in airway mucus. Sci Rep 5, 14644.

Giraud, C., Bernard, C.S., Calderon, V., Yang, L., Filloux, A., Molin, S., Fichant, G., Bordi, C., and de Bentzmann, S. (2011). The PprA-PprB two-component system activates CupE, the first non-archetypal Pseudomonas aeruginosa chaperone-usher pathway system assembling fimbriae. Environ Microbiol 13, 666–683.

Gomez, N.O., Tetard, A., Ouerdane, L., Laffont, C., Brutesco, C., Ball, G., Lobinski, R., Denis, Y., Plesiat, P., Llanes, C., et al. (2020). Involvement of the Pseudomonas aeruginosa MexAB-OprM efflux pump in the secretion of the metallophore pseudopaline. Mol Microbiol.

Gomez, N.O., Tetard, A., Ouerdane, L., Laffont, C., Brutesco, C., Ball, G., Lobinski, R., Denis, Y., Plesiat, P., Llanes, C., et al. (2021). Involvement of the Pseudomonas aeruginosa MexAB-OprM efflux pump in the secretion of the metallophore pseudopaline. Mol Microbiol 115, 84–98.

Gray, R.D., Imrie, M., Boyd, A.C., Porteous, D., Innes, J.A., and Greening, A.P. (2010). Sputum and serum calprotectin are useful biomarkers during CF exacerbation. J Cyst Fibros 9, 193–198.

Hood, M.I., and Skaar, E.P. (2012). Nutritional immunity: transition metals at the pathogen-host interface. Nat Rev Microbiol 10, 525–537.

Kaniga, K., Delor, I., and Cornelis, G.R. (1991). A wide-host-range suicide vector for improving reverse genetics in gram-negative bacteria: inactivation of the blaA gene of Yersinia enterocolitica. Gene 109, 137–141.

Kehl-Fie, T.E., and Skaar, E.P. (2010). Nutritional immunity beyond iron: a role for manganese and zinc. Curr Opin Chem Biol 14, 218–224.

Koch, B., Jensen, L.E., and Nybroe, O. (2001). A panel of Tn7-based vectors for insertion of the gfp marker gene or for delivery of cloned DNA into Gram-negative bacteria at a neutral chromosomal site. J Microbiol Methods 45, 187–195.

Kordes, A., Preusse, M., Willger, S.D., Braubach, P., Jonigk, D., Haverich, A., Warnecke, G., and Haussler, S. (2019). Genetically diverse Pseudomonas aeruginosa populations display similar transcriptomic profiles in a cystic fibrosis explanted lung. Nat Commun 10, 3397.

Lhospice, S., Gomez, N.O., Ouerdane, L., Brutesco, C., Ghssein, G., Hajjar, C., Liratni, A., Wang, S., Richaud, P., Bleves, S., et al. (2017). Pseudomonas aeruginosa zinc uptake in chelating environment is primarily mediated by the metallophore pseudopaline. Sci Rep 7, 17132.

Liberati, N.T., Urbach, J.M., Miyata, S., Lee, D.G., Drenkard, E., Wu, G., Villanueva, J., Wei, T., and Ausubel, F.M. (2006). An ordered, nonredundant library of Pseudomonas aeruginosa strain PA14 transposon insertion mutants. Proc Natl Acad Sci U S A 103, 2833–2838.

MacGregor, G., Gray, R.D., Hilliard, T.N., Imrie, M., Boyd, A.C., Alton, E.W., Bush, A., Davies, J.C., Innes, J.A., Porteous, D.J., et al. (2008). Biomarkers for cystic fibrosis lung disease: application of SELDI-TOF mass spectrometry to BAL fluid. J Cyst Fibros 7, 352–358.

Marguerettaz, M., Dieppois, G., Que, Y.A., Ducret, V., Zuchuat, S., and Perron, K. (2014). Sputum containing zinc enhances carbapenem resistance, biofilm formation and virulence of Pseudomonas aeruginosa. Microb Pathog 77, 36–41.

Mastropasqua, M.C., D’Orazio, M., Cerasi, M., Pacello, F., Gismondi, A., Canini, A., Canuti, L., Consalvo, A., Ciavardelli, D., Chirullo, B., et al. (2017). Growth of Pseudomonas aeruginosa in zinc poor environments is promoted by a nicotianamine-related metallophore. Mol Microbiol.

Mastropasqua, M.C., Lamont, I., Martin, L.W., Reid, D.W., D’Orazio, M., and Battistoni, A. (2018). Efficient zinc uptake is critical for the ability of Pseudomonas aeruginosa to express virulence traits and colonize the human lung. J Trace Elem Med Biol 48, 74–80.

Mikhaylina, A., Ksibe, A.Z., Scanlan, D.J., and Blindauer, C.A. (2018). Bacterial zinc uptake regulator proteins and their regulons. Biochem Soc Trans 46, 983–1001.

Neff, S.L., Doing, G., Reiter, T., Hampton, T.H., Greene, C.S., and Hogan, D.A. (2024). Pseudomonas aeruginosa transcriptome analysis of metal restriction in ex vivo cystic fibrosis sputum. Microbiol Spectr 12, e0315723.

Nelson, C.E., Huang, W., Zygiel, E.M., Nolan, E.M., Kane, M.A., and Oglesby, A.G. (2021). The Human Innate Immune Protein Calprotectin Elicits a Multimetal Starvation Response in Pseudomonas aeruginosa. Microbiol Spectr 9, e0051921.

Nielsen, F.H. (2000). Evolutionary events culminating in specific minerals becoming essential for life. Eur J Nutr 39, 62–66.

Palmer, L.D., and Skaar, E.P. (2016). Transition Metals and Virulence in Bacteria. Annu Rev Genet 50, 67–91.

Patzer, S.I., and Hantke, K. (1998). The ZnuABC high-affinity zinc uptake system and its regulator Zur in Escherichia coli. Mol Microbiol 28, 1199–1210.

Pederick, V.G., Eijkelkamp, B.A., Begg, S.L., Ween, M.P., McAllister, L.J., Paton, J.C., and McDevitt, C.A. (2015). ZnuA and zinc homeostasis in Pseudomonas aeruginosa. Sci Rep 5, 13139.

Rossi, E., Falcone, M., Molin, S., and Johansen, H.K. (2018). High-resolution in situ transcriptomics of Pseudomonas aeruginosa unveils genotype independent patho-phenotypes in cystic fibrosis lungs. Nat Commun 9, 3459.

Schalk, I.J., Abdallah, M.A., and Pattus, F. (2002). Recycling of pyoverdin on the FpvA receptor after ferric pyoverdin uptake and dissociation in Pseudomonas aeruginosa. Biochemistry 41, 1663–1671.

Schindelin, J., Arganda-Carreras, I., Frise, E., Kaynig, V., Longair, M., Pietzsch, T., Preibisch, S., Rueden, C., Saalfeld, S., Schmid, B., et al. (2012). Fiji: an open-source platform for biological-image analysis. Nat Methods 9, 676–682.

Smith, S., Rowbotham, N.J., and Charbek, E. (2022). Inhaled antibiotics for pulmonary exacerbations in cystic fibrosis. Cochrane Database Syst Rev 8, CD008319.

Son, M.S., Matthews, W.J., Jr., Kang, Y., Nguyen, D.T., and Hoang, T.T. (2007). In vivo evidence of Pseudomonas aeruginosa nutrient acquisition and pathogenesis in the lungs of cystic fibrosis patients. Infect Immun 75, 5313–5324.

Tuon, F.F., Dantas, L.R., Suss, P.H., and Tasca Ribeiro, V.S. (2022). Pathogenesis of the Pseudomonas aeruginosa Biofilm: A Review. Pathogens 11.

Turner, K.H., Wessel, A.K., Palmer, G.C., Murray, J.L., and Whiteley, M. (2015). Essential genome of Pseudomonas aeruginosa in cystic fibrosis sputum. Proc Natl Acad Sci U S A 112, 4110–4115.

Vermilyea, D.M., Crocker, A.W., Gifford, A.H., and Hogan, D.A. (2021). Calprotectin-Mediated Zinc Chelation Inhibits Pseudomonas aeruginosa Protease Activity in Cystic Fibrosis Sputum. J Bacteriol 203, e0010021.

Wakeman, C.A., Moore, J.L., Noto, M.J., Zhang, Y., Singleton, M.D., Prentice, B.M., Gilston, B.A., Doster, R.S., Gaddy, J.A., Chazin, W.J., et al. (2016). The innate immune protein calprotectin promotes Pseudomonas aeruginosa and Staphylococcus aureus interaction. Nat Commun 7, 11951.

Wang, S., Cheng, J., Niu, Y., Li, P., Zhang, X., and Lin, J. (2021). Strategies for Zinc Uptake in Pseudomonas aeruginosa at the Host-Pathogen Interface. Front Microbiol 12, 741873.

Wilson, G.B., Fudenberg, H.H., and Jahn, T.L. (1975). Studies on cystic fibrosis using isoelectric focusing. I. An assay for detection of cystic fibrosis homozygotes and heterozygote carriers from serum. Pediatr Res 9, 635–640.

Zhang, J., Zhao, T., Yang, R., Siridechakorn, I., Wang, S., Guo, Q., Bai, Y., Shen, H.C., and Lei, X. (2019). De novo synthesis, structural assignment and biological evaluation of pseudopaline, a metallophore produced by Pseudomonas aeruginosa. Chem Sci 10, 6635–6641.

