## Supplementary figures and images for "Pseudopaline-mediated zinc uptake by *Pseudomonas aeruginosa* determines specific clinically relevant phenotypes and infection outcome"

### Figure Sup

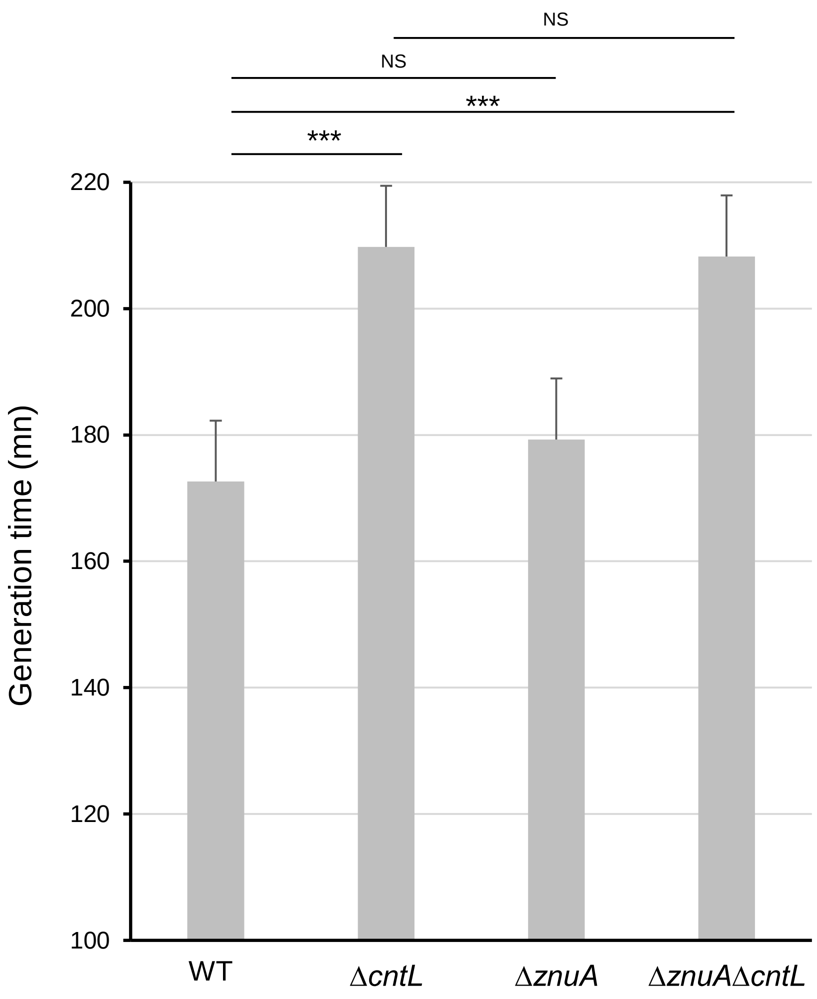

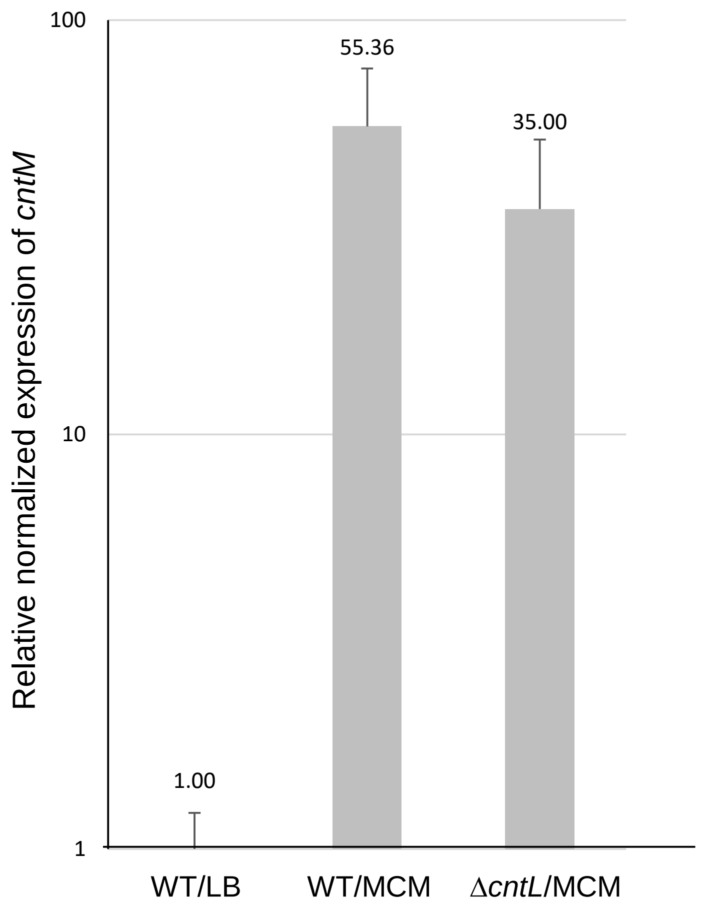

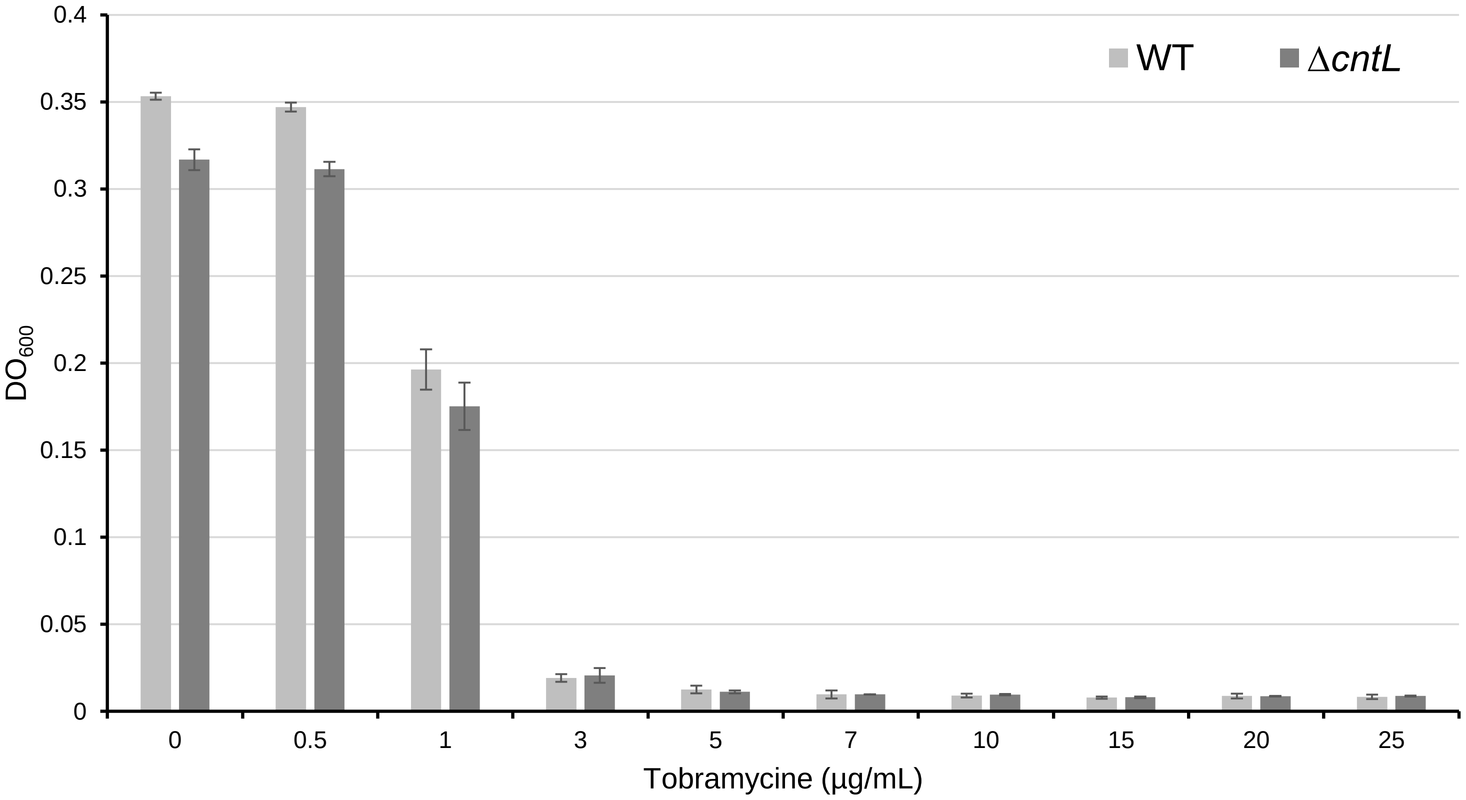

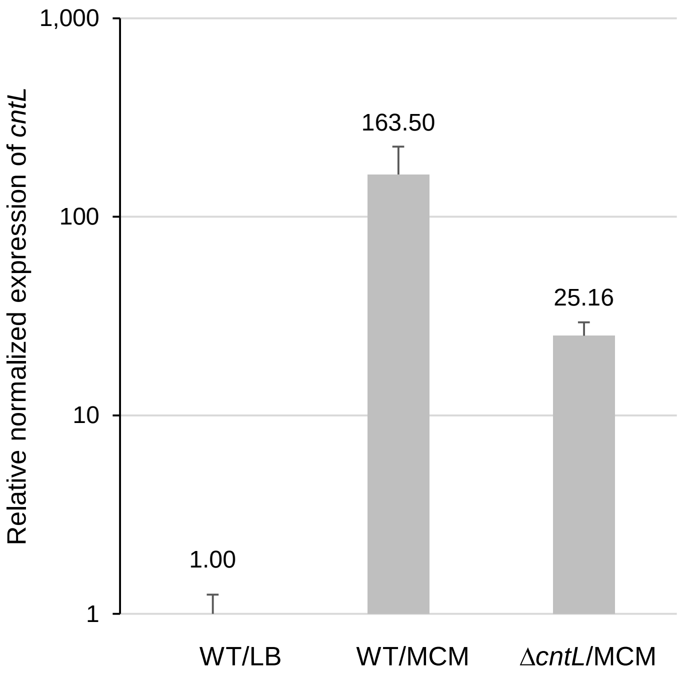
